# Effect of heterospecific and conspecific competition on individual differences in tadpole behaviour

**DOI:** 10.1101/2022.06.23.497343

**Authors:** Cammy Beyts, Maddalena Cella, Nick Colegrave, Roger Downie, Julien G. A. Martin, Patrick Walsh

## Abstract

Repeated social interactions with conspecifics and/or heterospecifics during early development may drive the differentiation of behaviour among individuals. This behavioural differentiation may occur through individuals behaving more different from each other on average and/or individuals behaving more consistently. Competition is a major form of social interaction and its impacts can depend on whether interactions occur between conspecifics or heterospecifics and the directionality of a response could be specific to different behavioural traits. To test this, we reared tungara frog tadpoles (*Engystomops pustulosus*) either in isolation, with a conspecific tadpole or with an aggressive heterospecific tadpole, the whistling frog tadpole, *Leptodactylus fuscus*. In each treatment, we measured the body size, activity, exploration and risk taking in the presence of a predator in focal *E. pustulosus* tadpoles six times during development. We used univariate and multivariate hierarchical mixed effect models to investigate the effect of treatment on mean behaviour and on among individual variance between and within individuals across behavioural traits. There was a strong effect of competition on behaviour, with different population and individual level responses across social treatments. Within their home tank, individuals were more consistent in their movements under conspecific competition but heterospecific competition caused more variance in the average movement among individuals. Behavioural responses were also trait specific as conspecific competition caused greater variability in movements among individuals in a novel environment. The results highlight that the impact of competition on inter-individual differences in behaviour is dependent on competitor species identity and is trait specific.

## Introduction

Among-individual variation in the behaviour of animals is well characterised and concerns among individual level variation in the mean behaviour of single traits and among individual level correlations between multiple traits (Sih et al. 2004; Réale et al. 2010; Dingemanse and Dochtermann 2013). Individual level differences in behaviour are thought to be maintained by genetic (co)variation and equal fitness payoffs associated with different behavioural strategies (Stamps 2007; Wolf et al. 2008; Mathot et al. 2012). Similarly, the niche specialisation hypothesis uses comparable statistical and biological concepts to understand how conspecific and heterospecific competition for food and space may drive among individual level differences in dietary preference, to allow limited resources to be partitioned among individuals (Bolnick et al. 2003; Araújo et al. 2011). These behavioural and ecological frameworks are now becoming integrated through the social (Bergmüller and Taborsky 2010; Montiglio et al. 2013) and behavioural niche (Kent and Sherry 2020) hypotheses which predict that conspecific and heterospecific competition will increase individual level differentiation in behaviour to reduce conflict over food and space. These multi-species interactions are important for understanding the proximate causes of inter-individual (co)variation in behaviour as well as individual level interactions which promote the coexistence of conspecifics at high density and co-occurrence of multiple species with similar resource needs (Bolnick et al. 2003; Briffa and Sneddon 2016; Kent and Sherry 2020; Sherry et al. 2020).

In the context of competition, there may be two ways in which individual level changes in behaviour may be partitioned (Figure 1). Individuals may diverge in their average behaviour, so that a broader range of behavioural strategies can be used to acquire a more diverse set of resources (Figure 1a; Prati et al., 2021; Preisser, Bolnick, & Grabowski, 2009). For example, less competitive individuals may be forced to forage at less optimal times of day or in less profitable foraging locations (Rychlik 2005; Frere et al. 2008; Harrington et al. 2009; Wauters et al. 2019). This would be detectable as an increase in the variance among individuals as individuals diverge in their average behaviour.

**Figure 1.**
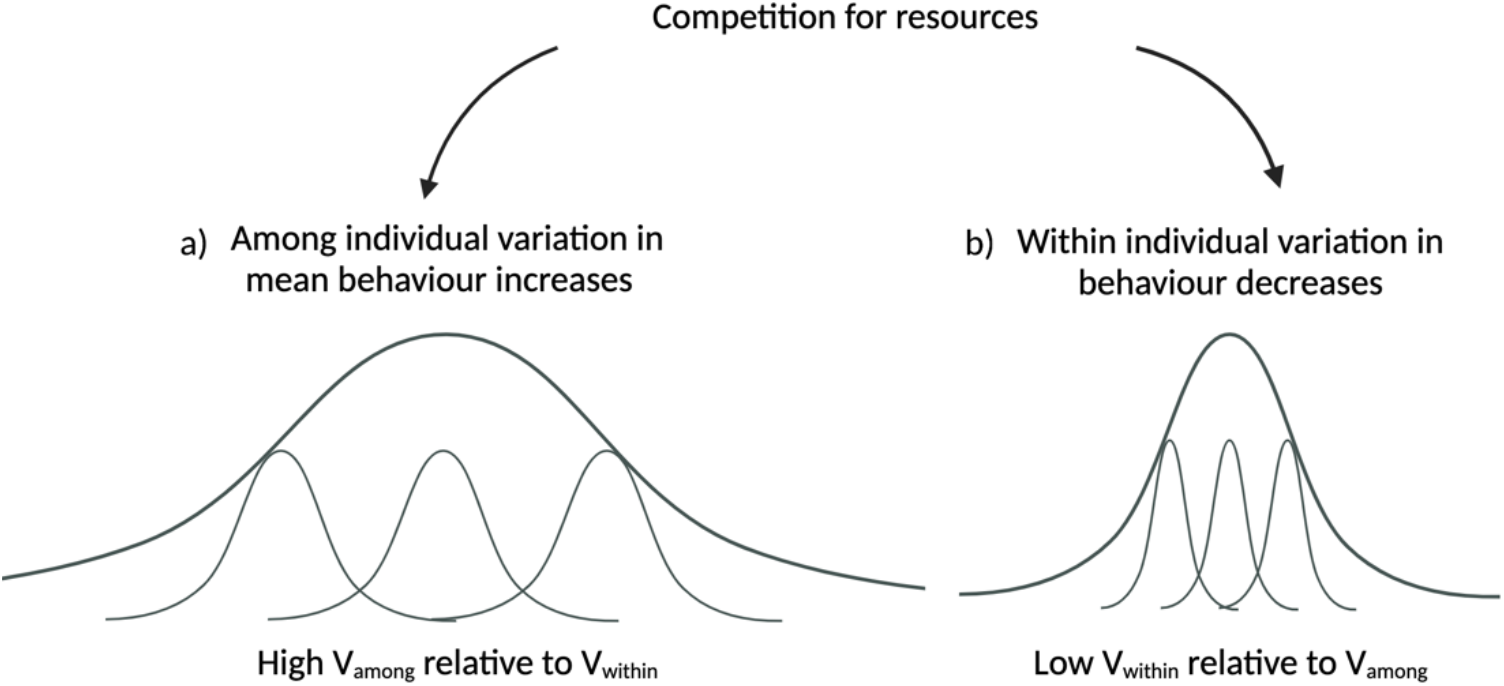
Conceptual illustration of two potential effects on individual level differences in behaviour in response to competition. The thin lighter lines are the behavioural responses of individuals and the darker thicker lines are the behavioural response of the population. The repeatability (R = V_among_ / V_among_ + V_within_) of plots a) and b) is the same. In plot a) individuals diverge in their behavioural strategy via an increase in among individual variation in mean behaviour. In plot b) individuals diverge in their behavioural strategy via a decrease in within individual variation.

Alternatively, competition for resources may also affect how consistent individuals are in their behavioural strategy, by influencing how variable individuals are within themselves (Stamps et al. 2012; Westneat et al. 2015). Here individuals may specialise in a particular microhabitat by showing greater consistency in their foraging behaviour(Figure 1b; Beaulieu & Sockman, 2012; Newell et al., 2014; Sherry et al., 2020). For example, each individual may specialise in foraging at a specific time of day within the most optimal foraging hours for that species. Here, individuals would diverge in their behaviour as they become less variable within themselves (Dingemanse and Dochtermann 2013) and would be detectable as a decrease in the variance within individuals. Consequently, under competition we may expect inter-individual level variation in behaviour to increase but the processes driving this change may be due to either an increase in among individual variance or a decrease in within individual variance. The repeatability statistic can be used to understand when competition may be driving individual level variation and the individual level variance components used to calculate repeatability can be used to determine whether it is variability among or within individuals which is responsible for this change (Bell et al. 2009; Nakagawa and Schielzeth 2010; Jäger et al. 2019). Repeatability will be high when variance among individuals is high relative to within individual variance or when within individual variance is low relative to among individual variance (Nakagawa and Schielzeth 2010; Dochtermann and Royauté 2019).

An individual’s behavioural strategy may also change across contexts (Stamps and Groothuis 2010; Arvidsson et al. 2017; Mitchell and Houslay 2021). This is because novel and risky contexts may result in potentially bolder or more cautious behaviours compared to familiar, low risk contexts (Carter et al. 2013; Perals et al. 2017; Kelleher et al. 2018). An individual’s perception of risk may further be dependent on the level and type of competition they are exposed to during development (Urszán et al. 2015; Han and Dingemanse 2017; He et al. 2017). Increased competition for resources may mean that individuals which are in greater need of resources may be prepared to take more risks and travel further distances in unfamiliar contexts (McNamara and Houston 1987; McNamara and Houston 1994; Anholt and Werner 1995; Anholt and Werner 1998). Therefore, patterns of inter-individual behavioural variation may be both influenced by the competitive environment as well as be context specific.

Different competitive environments may also favour specific combinations of behaviour traits (Bell 2005; Dingemanse et al. 2007; Fischer et al. 2016). In the absence of competition, there may only be a weak association between an individual’s behaviour in familiar, novel or risky contexts (Bell and Sih 2007; Dingemanse et al. 2007). However, exposure to competition may require individuals to up or down regulate their foraging activity across a range of contexts to secure additional resources (Bergmüller and Taborsky 2010). Consequently, conspecific and heterospecific competition may cause individuals to change their behaviour across multiple traits which would be detectable as correlations between traits at the among individual level (Garamszegi and Herczeg 2012; Dingemanse and Dochtermann 2013).

In this study we investigate the effect of both conspecific and heterospecific competition in the tungara frog tadpole (*Engystomops pustulosus*) which can be frequently found inhabiting the same temporary pools with the whistling frog tadpole (*Leptodactylus fuscus*) in Trinidad (Downie and Nicholls 2004). Both species have a similar development time of three weeks and occupy a similar ecological niche, suggesting a high level of resource overlap (Murphy et al. 2018; Santana et al. 2019; Atencia et al. 2020). The superior competitive ability of *L. fuscus* is thought to be attributed to its larger starting size and higher activity rates (Downie and Nokhbatolfoghahai 2006; Downie et al. 2008). Amphibian larvae represent an ideal life stage and group of organisms in which to investigate the effects of competition on inter-individual differences in behaviour and individual level correlations between traits (Urszán et al. 2015; Urszán et al. 2018). Across a variety of species, tadpoles compete with both conspecifics and heterospecifics for access to resources to fuel fast growth and development prior to metamorphosis and many of these interactions involve asymmetrical competition between species (Werner 1992; Bardsley and Beebee 2001; Richter-Boix et al. 2004; Smith et al. 2004; Richter-Boix et al. 2007; Ramamonjisoa and Natuhara 2017).

The behavioural traits we investigated were the levels of activity in a familiar context, exploration of a novel context and the propensity to take risks under predatory threat. In the activity assay, we predicted that conspecific and heterospecific competition would increase the repeatability of swimming behaviours through an increase in among individual variance in mean behaviour and/or increase in within individual variance. We also predicted that the repeatability of behaviour would differ between the three assay types and that behavioural differentiation would be greater under heterospecific compared to conspecific competition. Finally, we predicted that competition with conspecifics and heterospecifics would lead to correlations between traits at the among individual level, which may not be present in the absence of competition.

## Methods

### Study species and collection sites

We collected a total of 31 *E. pustulosus* and 25 *L. fuscus* foam nests from Lopinot Village, Trinidad (DMS: 10°41’21.7”N, 61°19’26.9”W) between June and July 2019, across four separate collection trips (see experimental design). We collected nests from pools located along a 400-meter length of road where both species are known to co-occur (Downie 2004; Figure 2). We placed each *E. pustulosus* nest into a separate container (dimensions: 145 x 100 x 55mm) containing water from the collection site and each *L. fuscus* nest into containers lined with a damp paper towel. As *L. fuscus* tadpoles rely on heavy rainfall to be washed into larval pool, they can suspend their development after hatching in the absence of water (Downie 1984; Downie 1994). However once submerged in water, their development continues as normal. Consequently, eight of the 25 *L. fuscus* nests collected had already hatched but had not developed beyond Gosner stage 27-28 (Gosner 1960).

**Figure 2.**
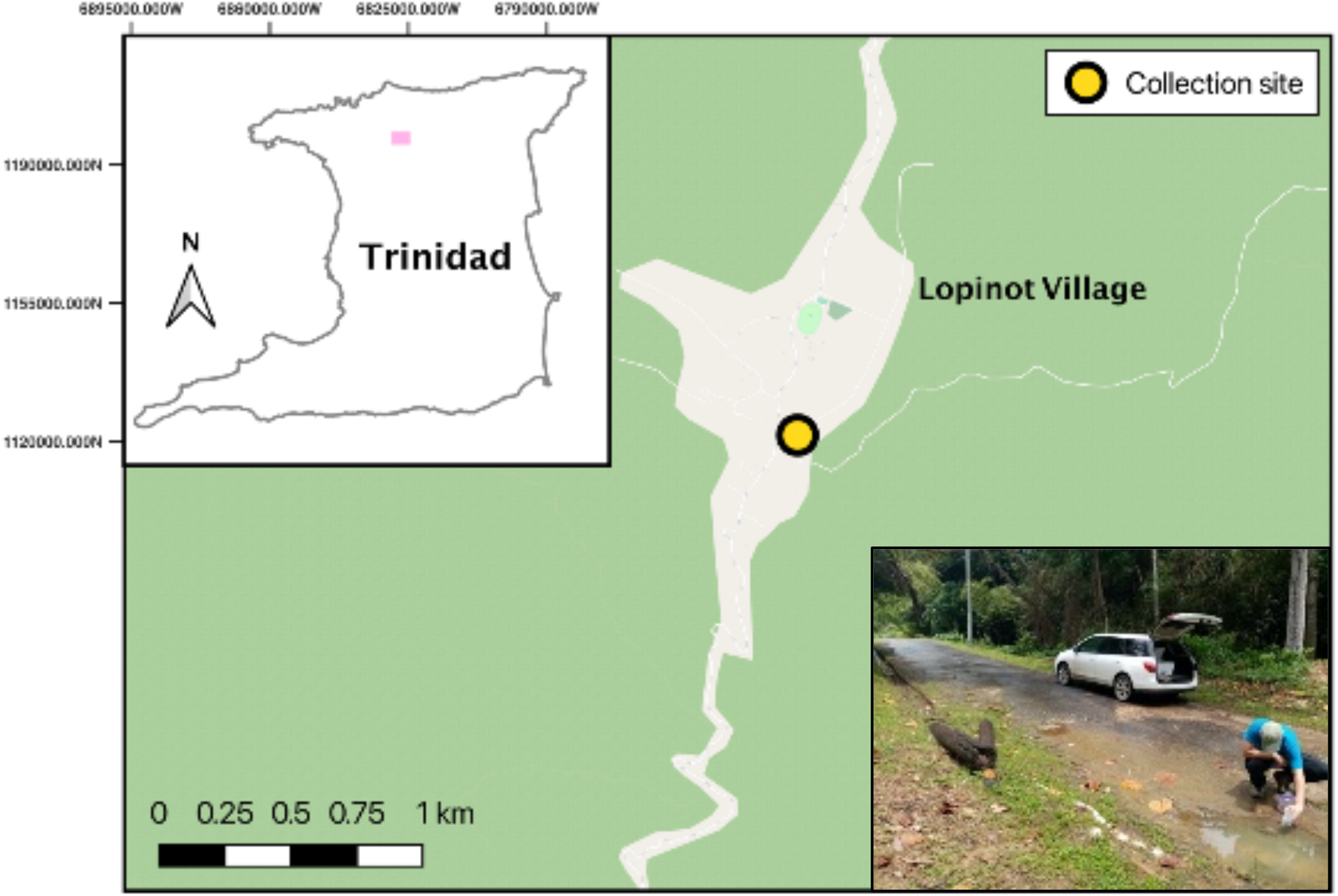
Map displaying the collection site of *Engystomops pustulosus* and *Leptodactylus fuscus* foam nests in Lopinot Village, Trinidad. A picture of the collection site is pictured in the bottom right panel. Image credit: Cammy Beyts

We transported the nests back to the William Beebe Tropical Research Station, “Simla”, (DMS10°41’30.7”N 61°17’26.4”W) located in Trinidad’s Northern Range, within two hours of collection. *E. pustulosus* tadpoles emerged from their eggs between 24 and 48 hours after collection. *L. fuscus* tadpoles from nests which had not already hatched were more variable in their emergence time, emerging between 24 to 96 hours post collection. We exposed nests and tadpoles to a 12.5L: 11.5D photoperiod and ambient temperatures ranging between 23.4 °C and 27.9°C (24.8°C ± 0.02 SD).

### Experimental design

We established three treatment groups to examine the impact of conspecific and heterospecific competition on *E. pustulosus* tadpole behaviour: 1) The “heterospecific treatment” contained one *E. pustulosus* tadpole and one *L. fuscus* tadpole (Figure 3A); 2) the “conspecific treatment” contained two *E. pustulosus* tadpoles, each from different nests (Figure 3B) and 3) the “no competition treatment” which contained an *E. pustulosus* tadpole housed in isolation (Figure 3C).

**Figure 3.**
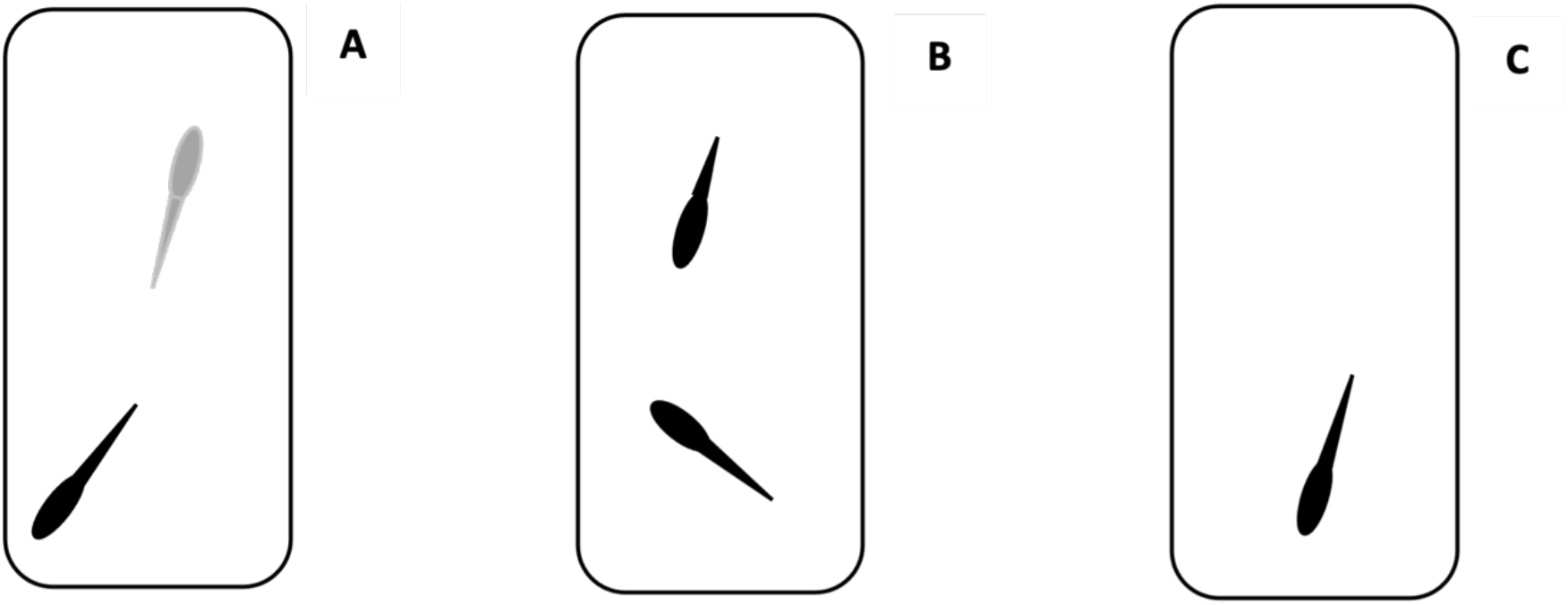
Illustration of treatment regimes. A: heterospecific treatment containing one *E. pustulosus* (black) and one *L. fuscus* (grey) tadpole. B: conspecific treatment containing one focal (full tail) and non-focal (shortened tail) *E. pustulosus* tadpoles. C: no competition treatment containing a solitary *E. pustulosus* tadpole.

We repeated the experiment over four consecutive batches, corresponding to the four collection trips. In each batch, we collected between 3 and 10 *E. pustulosus* foam nests 3 to 4 days before assigning tadpoles to their experimental treatments. *L. fuscus* nests were collected slightly earlier, 5-6 days prior to treatment assignment, due to the longer development time of *L. fuscus* eggs.

Within each batch, we assigned 15 focal *E. pustulosus* tadpoles to each treatment group, which were chosen at random from two *E. pustulosus* foam nests which hatched on the same day. This was to ensure that focal *E. pustulosus* tadpoles were the same age across each of the three treatment groups. Within the conspecific treatment, tadpoles from the two *E. pustulosus* nests were assigned as the focal or non-focal tadpole. To avoid potential weaker competitive dynamics among related individuals (Pakkasmaa and Aikio 2003; Yu and Lambert 2017), we obtained the non-focal tadpole in the conspecific treatment from the other nest to ensure that competitors were not siblings. To distinguish focal tadpoles in the conspecific treatment, we removed 1/3 of non-focal tadpole’s tail under MS-222 anaesthesia (Segev et al. 2015; Clarke et al. 2019). This distinguishing feature quickly disappeared, due to tail regeneration, so we subsequently distinguished focal tadpoles by visual differences in snout-vent-lengths that became apparent 3-4 days after the treatment commenced.

Focal and non-focal *E. pustulosus* tadpoles were at Gosner stage 25-26 when they were added to their treatment groups. *L. fucus* were more developed (Gosner stage 27-28) than the *E. pustulosus* tadpole in the heterospecific treatment, reflecting the natural circumstances of *L. fuscus* tadpoles in the wild typically entering breeding pools at a later stage of development (Downie 1984; Downie and Nicholls 2004).

We housed tadpoles in all treatments in plastic tanks (dimensions: 100 x 65 x 37mm), filled with 150ml of de-chlorinated, aerated tap water. We covered the tank sides in opaque tape, so tadpoles were not influenced by visual cues from tadpoles in adjacent tanks. We fed each tadpole in batches two through four with 7mg of ground fish food (TetraMin Tropical Fish Food Flakes) per day in the first week and 10mg in the second week. Due to a smaller initial size, we fed tadpoles in batch one with 3mg of food in the first week and 7mg in the second week. We left tadpoles undisturbed for five days following their assignment to treatments to allow them to acclimate and develop under their new social environment before starting behavioural assays. The experiment took 15 days from *E. pustulosus* hatching to the completion of the behavioural assays (Figure 4). This represents 60% of the larval period under ideal growth conditions. Tadpoles were returned to their sites of origin within 7 days of completing their final behavioural assay.

**Figure 4.**
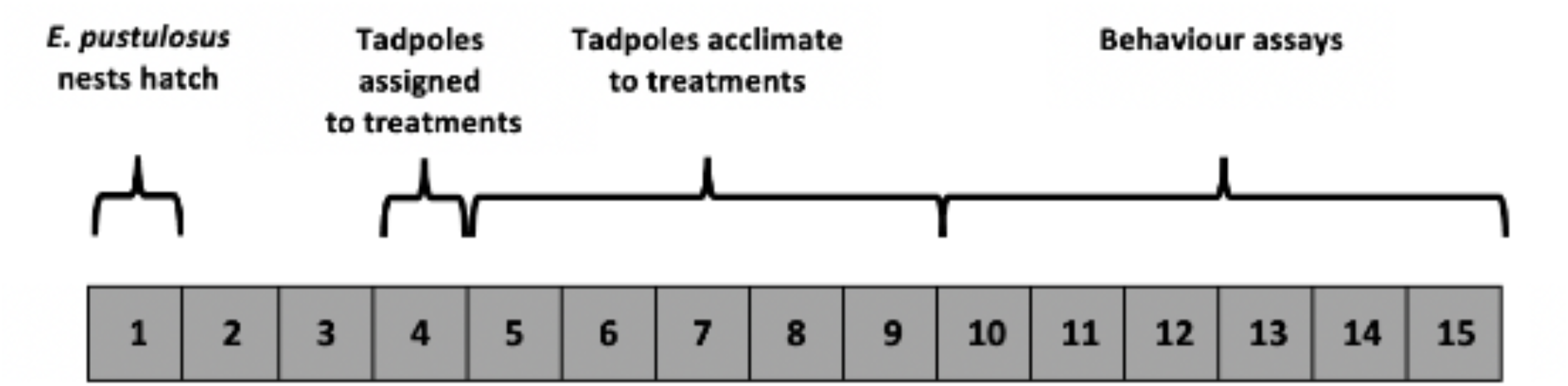
Summary timeline for an experimental batch. Numbers represent 24-hour days.

Across the four experimental batches, we collected data from 54 tadpoles in the no competition treatment, 56 focal tadpoles in the conspecific treatment and 51 tadpoles in the heterospecific treatment. 40 tadpoles across the three competition treatments died during the experiment which we did not include in the final tadpole count (supplementary material). We returned unused tadpoles and nests to their sites of origin within 7 days of collection.

### Behavioural assays

We recorded the behaviour of each tadpole in three behavioural assays, assessing activity, exploration, and predator risk-taking behaviour. We recorded each individual’s behaviour on six separate occasions over six consecutive days for each of the three assay types. We recorded assays in the same order (activity, exploration and predatory risk-taking) to limit the carry over effects of the more disruptive exploration and predatory risk-taking assays (Bell 2013). There was a total of 960, 993 and 909 trials recorded from the no competition, conspecific and heterospecific treatments respectively. We removed partial recordings from tadpoles that died before completing all 6 trials and recording errors (e.g., due to power outages) which were identified in 36/2898 trials.

We recorded all assays using one of four Canon Legria HF R86 camcorders, which we fixed in position (height: 450mm) above the activity tanks and exploration/predatory risk-taking arenas. We could film two tadpoles in separate, adjacent tanks/arenas simultaneously under one camera. The tanks/arenas could be positioned and removed from under the camera but were held in a fixed position during trials to assist with automated tracking software (see video processing). We filmed all the assays in a room adjacent to the laboratory where we performed husbandry procedures, under the same temperature and lighting conditions, to ensure that tadpoles would be undisturbed during filming.

### Activity assay

To measure activity levels in a familiar environment, we filmed the movement of focal *E. pustulosus* tadpoles in their home/rearing tanks over a 10-minute period. In the heterospecific and competition treatments, we removed non-focal tadpoles and placed them in a small cup of water from their home tank prior to starting the assay. All tadpoles were left undisturbed for 10 minutes prior to filming to allow them to acclimatise.

### Exploration assay

To quantify individual exploration of a novel environment, we filmed focal tadpole movements in a novel arena (dimensions: 29.8 x 19.5 x 4.9 mm; iDesign, UK), filled with 500ml of aerated tap water and warmed to lab temperature. The arena consisted of an acclimation zone (AZ) which opened to a central corridor with four compartments on both the left- and right-hand sides (Figure 5). To start a trial, we transferred one focal tadpole to the AZ and left them to acclimate for 10 minutes. We covered the top of the AZ with an opaque barrier to prevent disturbance from the investigator, and during acclimation we sealed the entrance to the corridor with an opaque removeable barrier. After acclimation, the investigator lifted the front portion of the barrier (the top barrier remained in position), providing the tadpole with access to the arena, and the tadpole’s movements were recorded over 15 minutes. The arena was cleaned between trials using tap water and fresh water was used for each new trial and tadpole.

**Figure 5.**
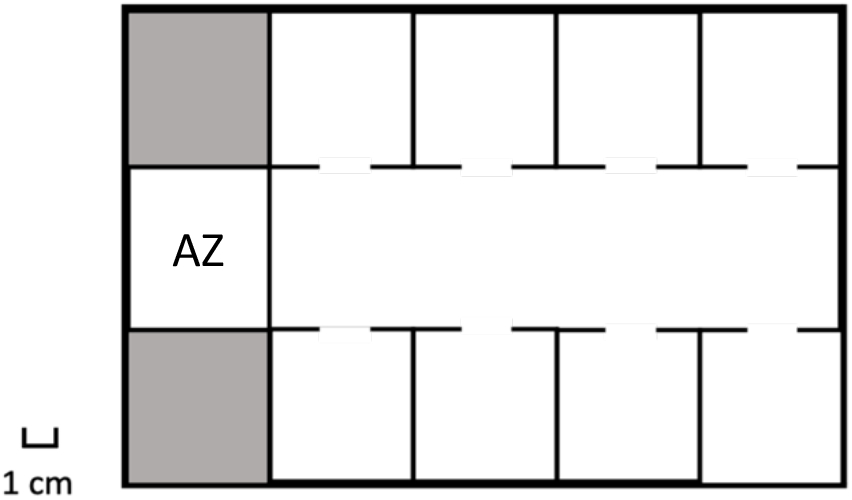
Diagram of the exploration arena. Tadpoles started the trial in the acclimation zone (AZ), spending 10 minutes behind an opaque barrier. After acclimation, tadpoles were free to explore the central corridor zones and adjacent zones to the left and right. Shaded areas represent unused sealed zones.

### Predator risk-taking assay

To quantify individual exploration of a new environment under predation risk, we recorded tadpole movements in the presence of visual and olfactory cues from a dragonfly larvae predator (family: Gomphidae) in a novel arena (dimensions: 17 x 12.5 x 4.6; Western Boxes, UK). Each arena consisted of a covered acclimation zone (AZ), an open zone (OZ) in which the tadpole could explore and a predator zone (PZ) which allowed visual cues and olfactory cues of the predator to pass through to the OZ (Figure 6). To start a trial, a dragonfly larva was placed into the arena PZ and the focal tadpole was placed into the arena AZ. The AZ was sealed with an opaque barrier to allow tadpoles to acclimate. 10ml of predator conditioned water was also added into the OZ to act as an additional predator olfactory cue after tadpoles and predators were added to the AZ and PZ respectively. Tadpoles were given 10 minutes to acclimate within the AZ before the barrier between the AZ and OZ was removed, we then recorded tadpole movements over 15 minutes. The arena was cleaned between trials using tap water and fresh water was used for each new trial and tadpole. Dragon fly larvae were collected from the Aripo Savannah in Trinidad, where both *E. pustulosus* and *L. fuscus* were also observed to co-occur alongside the dragonfly larvae. When not used in assays, we housed the dragonfly larvae in an 11L Perspex tank and fed them with four *E. pustulosus* tadpoles (which had died of natural causes) each morning.

**Figure 6.**
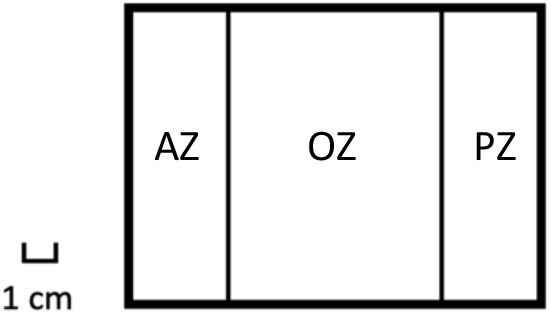
Diagram of the predation arena. AZ represents the acclimation zone where tadpoles acclimated to assay conditions. PZ represents the predator zone which contained a live dragonfly larval predator. OZ represents the open zone where the tadpole could explore when the opaque AZ barrier was removed. The barrier between the OZ and PZ was transparent and perforated to allow visual and olfactory predator cues to pass into the OZ.

### Video processing

Post filming, to reduce storage space and increase processing speed, all videos were re-sized to 640×360 pixels and the activity and exploration videos were reduced from 25fps to 1fps, using the command line tool ffmpeg (Tomar 2006). The predation assay trials were reduced to a higher frame rate of 5fps to capture the faster movements of tadpoles in this assay. In all three assays, we measured the total distance a tadpole travelled in pixels using a custom-written tracking tool (written by C.Beyts) developed in Python v3.0 and using the OpenCV v4.4 library. The tracking code can be found on Github (see data availability). In the exploration and predatory risk-taking assays, tadpoles that did not leave the acclimation zone received a distance score of 0.

### Morphological measures

The snout vent length (SVL) of each tadpole was measured in FIJI v2.0 (Schindelin et al. 2012) to the nearest 0.1mm from activity assay recordings as a measure of body size. Measurements were taken from each activity trial to give six SVL measurements for each tadpole.

All procedures were approved by the University of Edinburgh ethics committee, under the assessment pwalsh1-0001. Permits to collect *E. pustulosus* and *L. fuscus* were obtained from Trinidad’s Forestry and Wildlife Division.

## Statistical Analysis

We estimated the effect of treatment on tadpole body size and tadpole behaviour in two separate models using a Bayesian approach.

### Treatment effects on tadpole body size

To estimate the effect of treatment on tadpole body size we fitted a univariate linear mixed model with a Gaussian error distribution (Dingemanse and Dochtermann 2013). Body size was scaled prior to analysis. We included a fixed effect of treatment (no competition, conspecific and heterospecific treatment) to estimate the effect the social environment had on the scaled average body size of tadpoles. We also included a fixed effect of trial (fitted as a continuous covariate, coded from 0 to 5) to estimate how body size changed from trial one to six. We included a random effect of tadpole Egg Mass ID and Tadpole ID in the model.

### Treatment effects on tadpole behaviour

To estimate the effect of social treatment on i) the population mean behaviour, ii) variance among individuals and iii) variance within individuals, we fitted a multivariate generalized linear mixed model (Dingemanse and Dochtermann 2013) of total distance moved in the activity, exploration and predation risk-taking assays. The multivariate model allowed all parameters i-iii to be estimated for activity, exploration and predation risk-taking behaviours simultaneously as well as the pairwise correlations between parameter ii for each behavioural assay. Given that total distance moved is a variable constrained to be positive and can be bounded to zero in some assays, a log-normal distribution is appropriate. As there were a high proportion of exploration and predation-risk-taking trials where tadpoles never left the acclimatation zone (54% and 56% of trials respectively), we used a hurdle lognormal distribution for these assays. A hurdle log-normal distribution is in fact a mixture distribution combining a binomial process and a log-normal process. This is adequate for the exploration and predation-risk assays, where the tadpoles decide to leave the acclimation zone or not and then explore the arena. One of the advantages of the hurdle log-normal distribution is that it additionally allowed us to look at a final population level parameter as to whether tadpoles iv) left the acclimation zone or not. We used the log normal distribution for the activity data so that the distance measures for all three assays could be estimated on the same log scale and aid comparison of results between the three assays in the multivariate model.

The same fixed and random effect structure used in the body size univariate model was fitted to the activity, exploration and risk-taking assays in the multivariate model. We fitted two models, one where body size was fitted as a fixed effect and one where body size was not fitted to the model. However, the inclusion of body size did not change the study conclusions and thus was kept in the model. To estimate the effect of treatment on among and within individual variance in each assay, we allowed among-individual and residual variance to vary with each treatment. To estimate population differences in whether tadpoles left the acclimation zone or not for each treatment, we fitted a treatment specific fixed effect to the hurdle model for the exploration and predatory risk-taking behaviours.

Estimating among individual variance in mean behaviour in activity, exploration and predator risk-taking behaviours provided a 3×3 covariance matrix allowing us to estimate the correlations in mean behaviour between each assay for each of the three social treatments. Given that tadpoles were only exposed to one social treatment, we could not estimate the correlation across treatments at the individual level. We converted all covariance estimates into correlations to aid the interpretation of results.

All models were fitted using the brms package v2.15 (Bürkner 2018) within R v4.0 (RCoreTeam 2013). We used uninformative or weak priors on all parameters (Gelman et al. 2013) which included wide normal priors for fixed effects, Half-Student priors for variance parameters and LKJ correlation priors for correlations. The models met all assumptions on convergence and autocorrelation and posterior predictive checks were used to determine if the model fitted the observed data (Gelman et al. 2013).

## Data availability

The data and code for this study are available on GitHub https://github.com/cammybeyts/tungara_tadpole_competition

## Results

### The effect of treatment on body size

Tadpoles from the no competition treatment were larger than tadpoles from the heterospecific treatment and marginally larger than tadpoles in the conspecific treatment (Figure 7). Tadpoles experiencing conspecific competition were also larger than individuals experiencing heterospecific competition (Figure 7). There was among individual variance associated with Egg Mass ID and Tadpole identity (supplementary materials, Table S1). Tadpole body size increased from trial 1 through to trial 6 (Supplementary Materials, Table S1).

**Figure 7.**
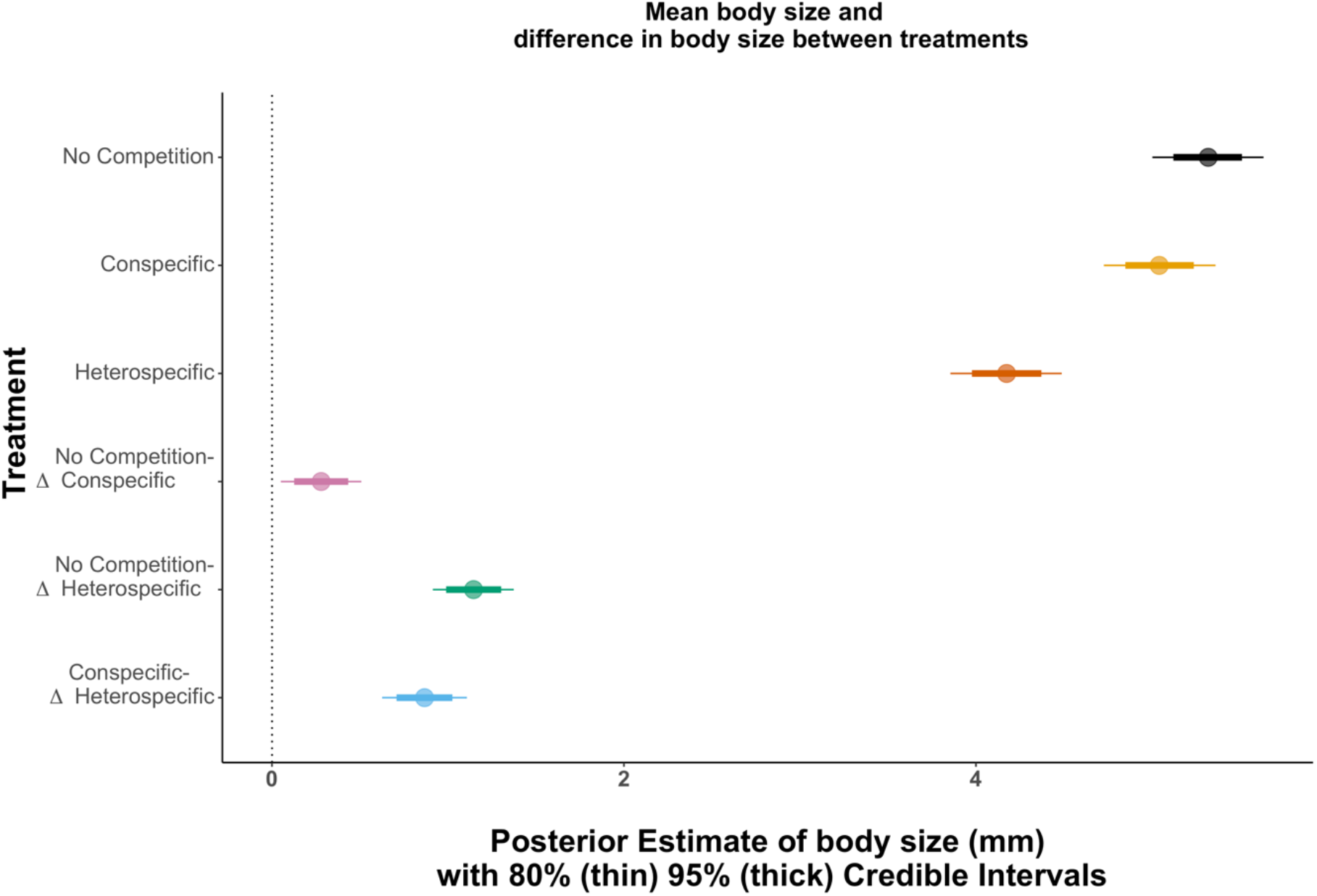
Treatment effects on tadpole overall mean body size. Points indicate posterior estimates for mean values and associated 80% (thick) and 95% (thin) credible intervals. Higher values indicate greater means. Estimates are displayed for No Competition (black), Conspecific (orange) and Heterospecific (red) treatment groups. The contrasts between treatments are displayed as the difference between No competition and Conspecific (purple), No Competition and Heterospecific (green) and Conspecific and Heterospecific (blue) treatment groups. Contrasts are displayed as absolute values.

### The effect of treatment on behaviour

At a population level, treatment did not affect the average distance tadpoles swam in the activity or predation risk-taking assays (Figure 8A). In the exploration assay, tadpoles in the no-competition treatment swam further than tadpoles exposed to competition, regardless of the type of competition (Figure 8A).

**Figure 8.**
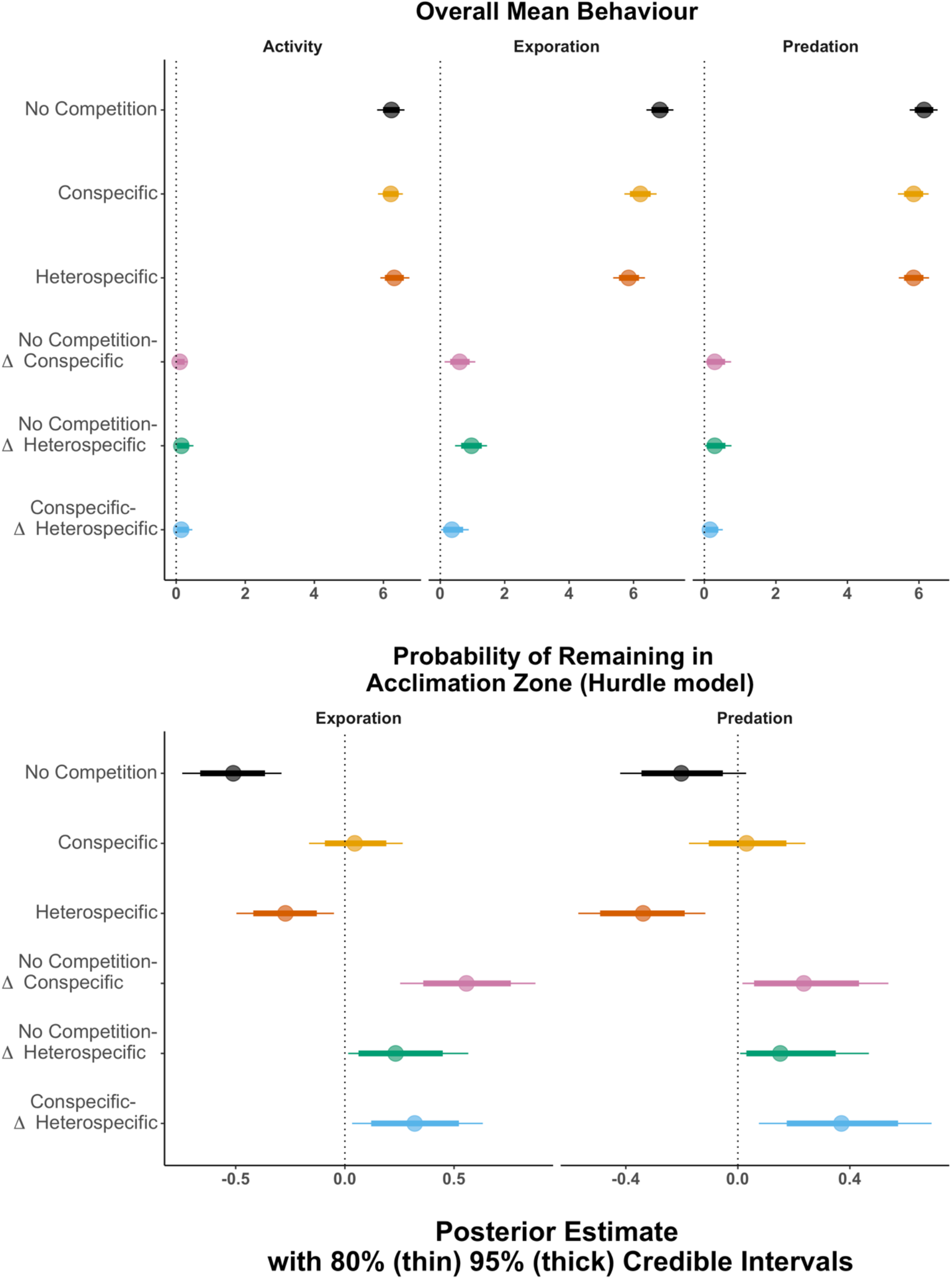
Treatment effects on (A) overall mean behaviour and (B) probability of remaining in the acclimation zone. Points indicate posterior estimates for mean values and associated 80% (thick) and 95% (thin) credible intervals. Higher values indicate greater means and higher probability of remaining in the acclimation zone. Estimates are displayed for No Competition (black), Conspecific (orange) and Heterospecific (red) treatment groups. The contrasts between treatments are displayed as the difference between No competition and Conspecific (purple), No Competition and Heterospecific (green) and Conspecific and Heterospecific (blue) treatment groups. Contrasts are displayed as absolute values.

In the exploration assay, tadpoles in the conspecific treatment were the most likely to remain in the acclimation zone compared to tadpoles in the no competition and heterospecific treatments (Figure 8B). The predation assay was similar, with tadpoles in the conspecific treatment being more likely to remain in the acclimation zone than in the heterospecific treatment (Figure 8B).

The among individual variance was non negligible in all 3 treatments for activity. Among individual variation in movements in the heterospecific treatment was larger than that observed in the no-competition treatment and marginally larger than the level observed in the conspecific treatment (Figure 9A). However conspecific competition caused greater variability in movements among individuals in a novel environment within the exploration assay. Among individual variance in movement levels did not change between treatments in the exploration assay (Figure 9B). Tadpoles did not show among individual variation in their mean levels of predatory risk-taking behaviours (Figure 9B, Supplementary Materials Table S2) and these did not change between treatments (Figure 9B).

**Figure 9.**
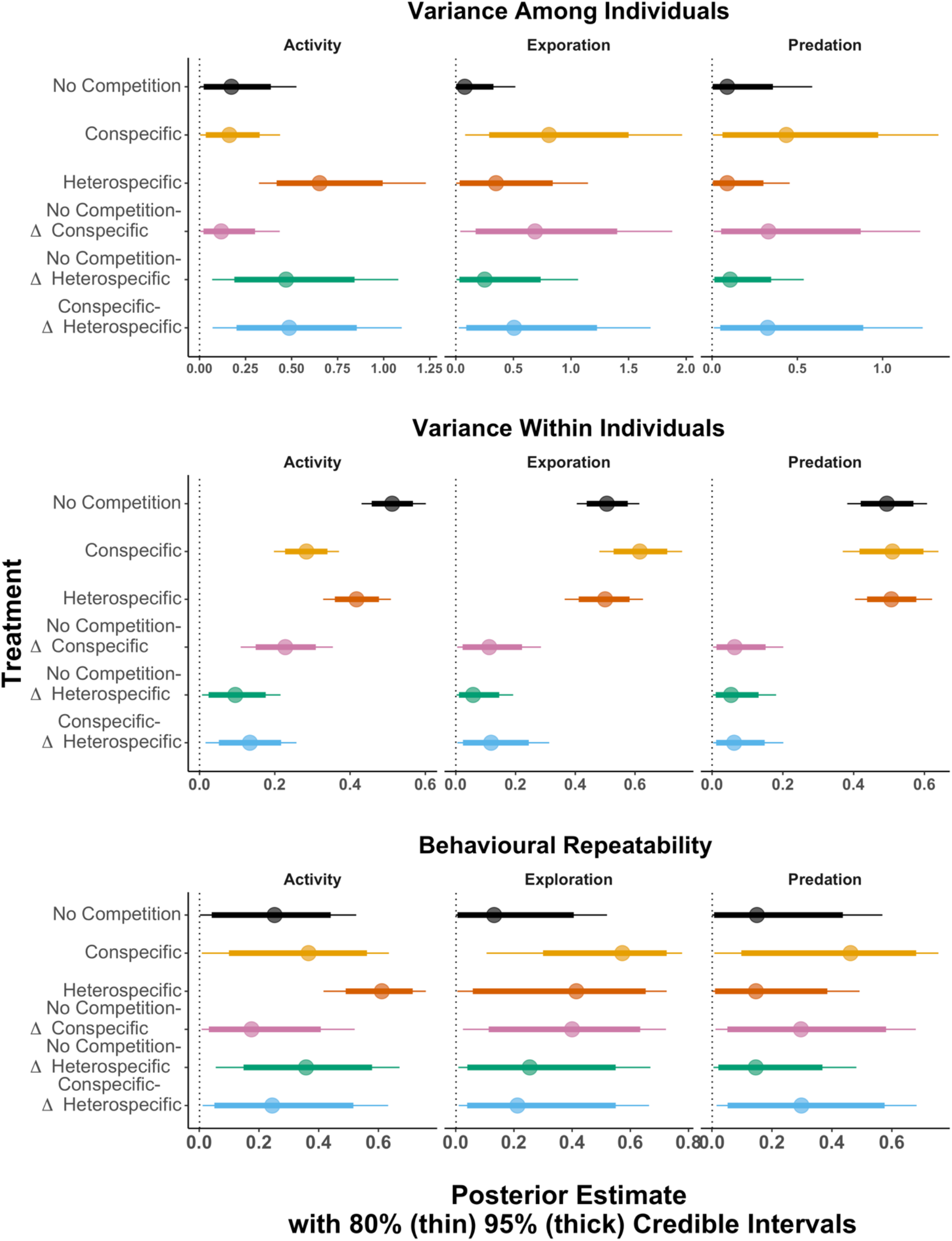
Treatment effects on (A) variance among individuals, (B) variance within individuals and C) behavioural repeatability for activity, exploration and predatory risk-taking behaviour. Points indicate posterior estimates for mean values and associated 80% (thick) and 95% (thin) credible intervals. Higher values indicate greater means and higher probability of remaining in the acclimation zone. Estimates are displayed for No Competition (black), Conspecific (orange) and Heterospecific (red) treatment groups. The contrasts between treatments are displayed as the difference between No competition and Conspecific (purple), No Competition and Heterospecific (green) and Conspecific and Heterospecific (blue) treatment groups. Contrasts are displayed as absolute values.

At the within individual (residual) level, tadpoles in the conspecific treatment were more consistent in their activity movements than tadpoles in the no competition and heterospecific treatments (Figure 9B, Supplementary Materials Table S2). Treatment did not affect the level of within individual variation in tadpoles in the exploration or predator risk-taking assays and the level of within individual variance did not differ between treatments for these assays (Figure 9B).

The activity levels of tadpoles were only repeatable within the heterospecific treatment and was greater than the level of repeatability observed within the no competition treatment (Figure 9C). Owing to an increase in among individual variance in mean exploration behaviour, tadpoles were more repeatable in their exploration behaviour within the conspecific treatment, but the level of repeatability did not change among treatments (Figure 9C). Tadpole behaviour was not repeatable in any treatment within the predation assay (Figure 9C).

There was no correlation in among individual variance in mean behaviour either within assays or between assays (Supplementary Materials Table S3). These correlations also did not differ between treatment regimes (Supplementary Materials Table S4).

## Discussion

Competition impacts inter-individual variation in tadpole behaviour with different patterns of among and within individual level variation observed in response to conspecific and heterospecific competitors. Within their home tank, heterospecific competition caused the repeatability of swimming movements to increase as individuals diverged in their swimming behaviour at the among individual level. In contrast, repeatability did not change under conspecific competition, but individuals became more consistent in their swimming movements. Behavioural responses were also trait specific as conspecific competition caused greater variability in movements among individuals in a novel environment. The results show that the impact of competition on inter-individual differences in behaviour is dependent on competitor species identity and is trait specific.

Ecological theory predicts that individuals can alleviate competition for resources through prey specialisation (MacArthur 1958; Bolnick et al. 2003). Changes in foraging behaviour may also be important in promoting co-existence where there is high resource overlap (Kent and Sherry 2020; Sherry et al. 2020). In our experiment, conspecific and heterospecific competition affected the among and within individual components of tadpole activity behaviour independently and this suggests that behavioural mechanisms for reducing conflict over contested resources may be different for single and multispecies interactions. In the heterospecific treatment, the biological mechanisms driving among individual patterns of activity behaviour may reflect differences in susceptibility of *E. pustulosus* individuals to competition owing to individual level differences in body size or basal metabolic rate (Careau et al. 2008; Biro and Stamps 2010; Kelleher et al. 2017). Consequently, the wider range of swimming movements among individuals may have reflected some individuals conserving energy by swimming shorter distances whilst others may have swum further to find more foraging opportunities. Alternatively, differences in morphology and/or behaviour among *L. fucus* individuals may have contributed to the diversity of behavioural responses observed in the focal *E. pustulosus* tadpoles via indirect effects (Wolf et al. 1998; Wilson et al. 2009; Jäger et al. 2019).

In the conspecific treatment, the increase in behavioural consistency without the corresponding change in behavioural repeatability suggests that individuals were not partitioning resources through behavioural specialisation. A more likely explanation is that the increased consistency in swimming movements was to allow focal tadpoles to converge their activity with the behaviour of the non-focal tadpole (Wolf et al. 2011). This may be beneficial for promoting increases in foraging gains through group foraging (Rook and Penning 1991; Rands et al. 2014), reduce the costs of locomotion (Marras et al. 2015) or provide increased protection from predators (Landeau and Terborgh 1986; Szulkin et al. 2006). In fish shoals, individuals may conform in their behaviour to produce coordinated changes in direction and bursts of speed (Jolles et al. 2018; Sankey et al. 2019; Jolles et al. 2020). This may be mediated by a decrease in behavioural variation both between each other and within themselves (Webster et al. 2007; Magnhagen and Bunnefeld 2009; Herbert-Read et al. 2013). Whilst shoaling behaviour has not been reported in *E. pustulosus* tadpoles, other larvae of anuran species such as cane toads (*Rhinella marina*) and common toads (*Bufo bufo*) are known to form dense aggregations (Wassersug et al. 1981; Griffiths and Foster 1998) where behavioural conformity may be important. To elucidate whether the decrease in within individual variance in response to conspecifics was driven by competition over resources or behavioural conformity, future studies could record the behaviour of both focal and non-focal individuals. If both individuals show similar patterns of behaviour and low within individual variance, this would indicate behavioural conformity over behavioural specialisation.

In addition to the species identity of a competitor impacting the stability or individuality of behaviour, we found the effect of competitive treatment was highly trait dependent. In particular, the level of within and among individual variation in swimming movements in the home tanks had no relation to the level of inter-individual variation within a novel or high predation risk context. Consequently, studies which only consider individual level behavioural responses in a single trait are likely to miss elements of behavioural variation that could be relevant in other ecological contexts. There was also no evidence that competition could alter the structure of behavioural syndromes within a population.

In the conspecific treatment, whilst tadpole movements were highly consistent in their home tanks, individuals showed variance in their mean exploration levels. As novel environments may present new foraging opportunities, consistent differences in state among focal individuals may have caused low state individuals to take greater foraging risks to secure additional feeding opportunities (Rands et al. 2003; Montiglio et al. 2013). Differences in mean exploration behaviour among individuals may also represent different foraging strategies, but these may be constrained in home tanks when individuals have repeatedly interacted with conspecifics during development. The lack of individual level variation under predation risk may further highlight that the high risk predators pose may limit the scope for multiple strategies, regardless of an individual’s prior exposure to competition.

Since low numbers of tadpoles left the acclimation zone, our power to detect differences in the repeatability of behaviour between treatments was limited (Martin et al. 2011; Dingemanse and Dochtermann 2013). Nevertheless, tadpoles in the conspecific treatment were the least likely to leave the acclimation zone across both the exploration and predation assays compared to tadpoles housed in isolation or with a heterospecific. This provides further support for our findings that behavioural responses to competition were both trait specific and dependent on the species identity of the competitor. Jolles et al (2016) suggested that testing fish in isolation when they had previously been housed in groups may induce stress (Gallup and Suarez 1980) compared to individuals which had always been housed alone. This may contribute to a reduced propensity for individuals to take risks (Jolles et al. 2016). A similar mechanism could explain the high number of tadpoles in the conspecific treatment which remained in the acclimation zone in the present study, compared to the increased exploration levels in the no competition treatment.

## Conclusions

This study shows that both conspecific and heterospecific competition can impact inter-individual differences in behaviour but may be mediated through different behavioural mechanisms, affecting among and within sources of individual level variation independently. As highlighted by the effect of conspecific competition on activity and exploration behaviour, this study also demonstrates that responses to competition is trait dependent. Future investigations should consider how individual level variation in behaviour may change in response to early life conditions depending on the behavioural traits investigated.

## Acknowledgements

We would like to thank Professor Paul Hoskisson, Greig Muir and members of the 2019 Glasgow Trinidad Expedition for their help in collecting tadpoles and running behavioural assays in this study.

