## Supplemental data for "Effect of heterospecific and conspecific competition on individual differences in tadpole behaviour"

Supplementary materials

### 1. Comparison of tadpole mortality rates across treatments

In total, across the four experimental batches, we collected data from 54 tadpoles in the no competition treatment, 56 focal tadpoles in the conspecific treatment and 51 tadpoles in the heterospecific treatment. 40 tadpoles died during the experiment which are not included in the final tadpole count. This included 10 tadpoles in each of the no competition and conspecific treatments and 20 tadpoles in the heterospecific treatment. This highlighted the increased severity of the heterospecific treatment in comparison to the no-competition (three sample proportion test with Bonferroni-Holm adjustment: X-squared = 8.26, df = 2, *p* = 0.038) and conspecific (*p* = 0.038) treatment groups. There was no difference between the number of tadpoles which died in the no-competition and conspecific treatment groups (*p* = 0.50).

### 2. Effect of treatment on mean tadpole body size

Table S1. Posterior estimates for fixed and random effects and their 95% credible intervals (CI) for treatment effects on tadpole body size.

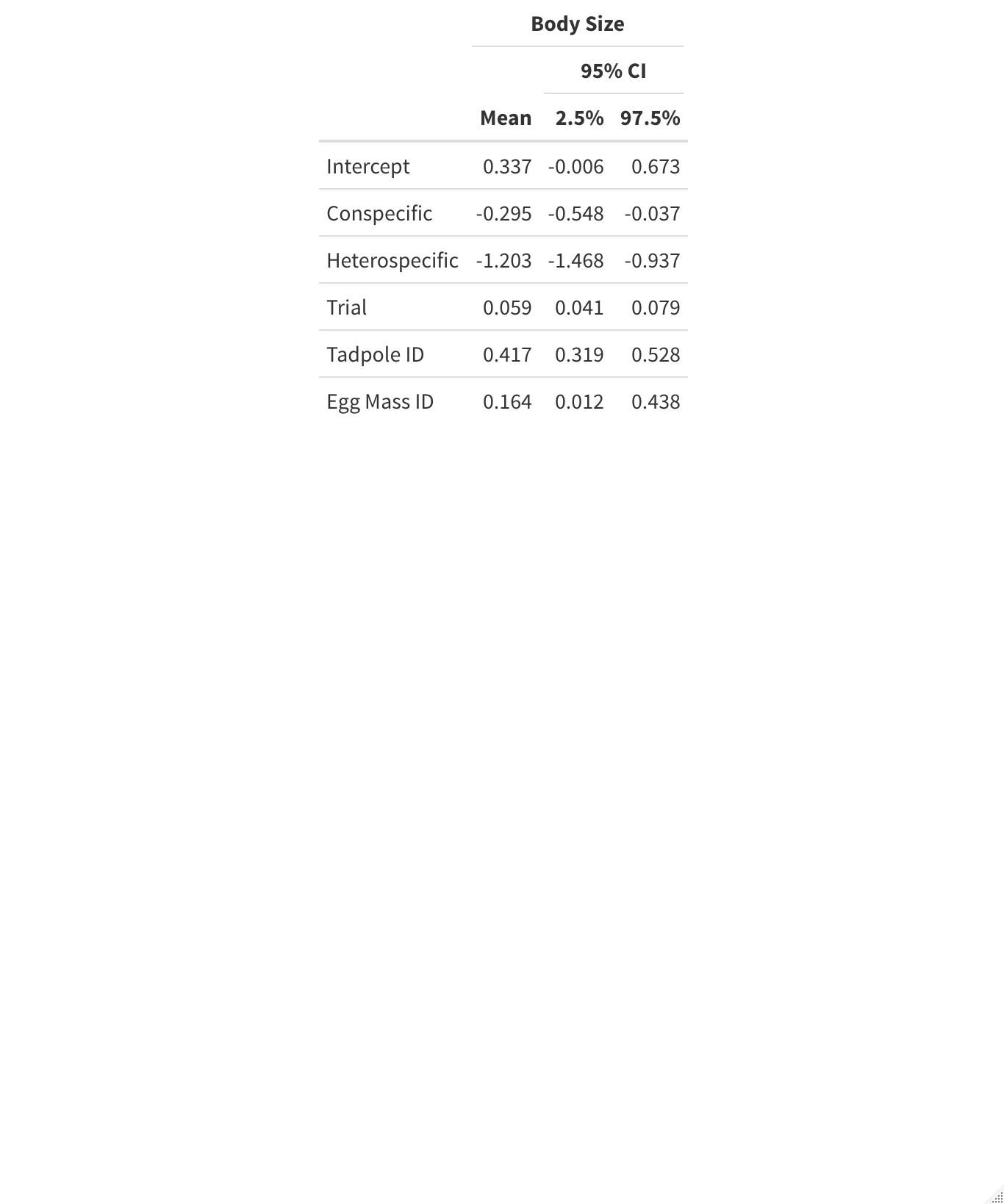

### 3. Effect of treatment on tadpole behaviour

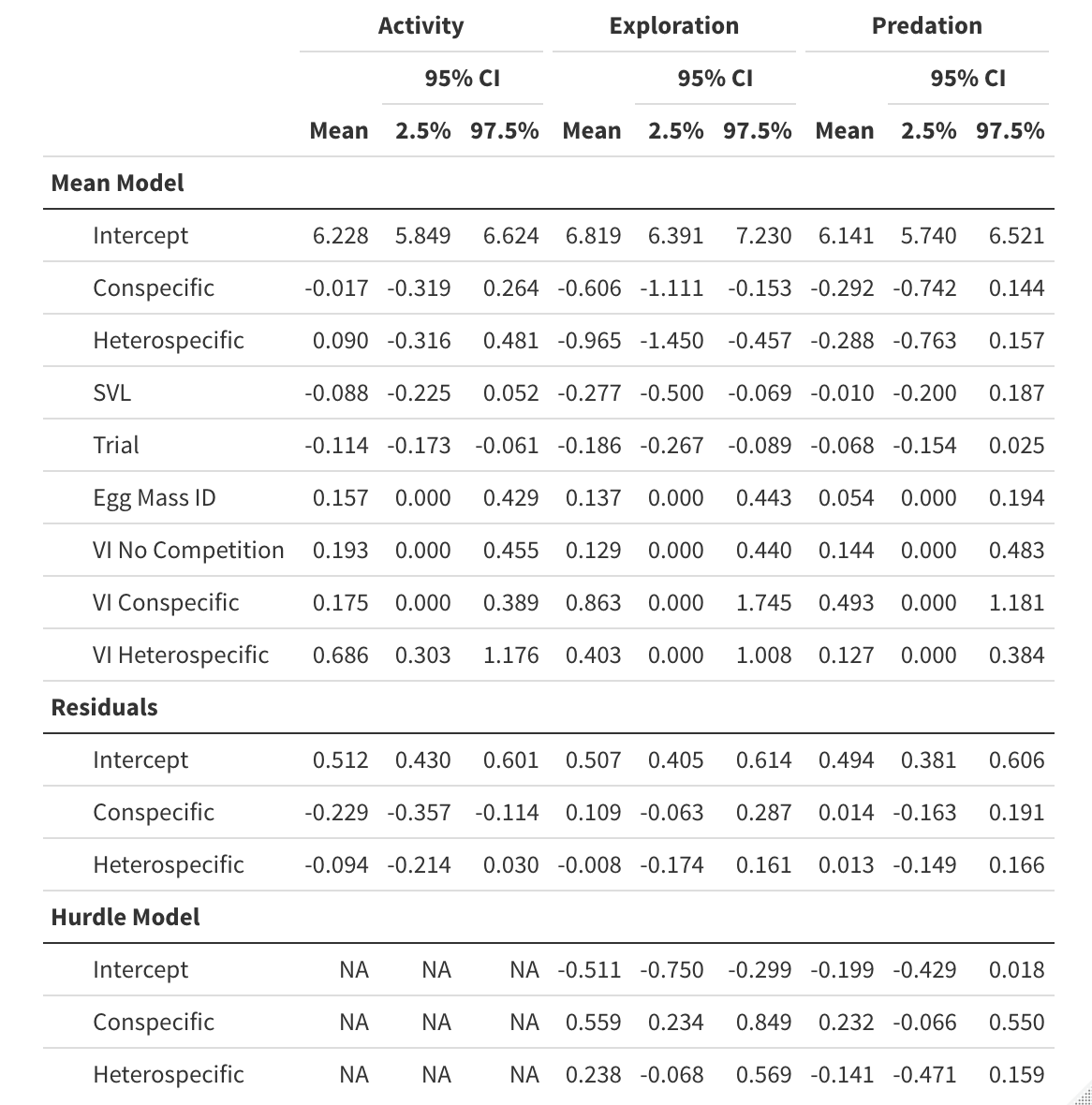

Table S2. Posterior estimates for fixed and random effects in the mean, residual and hurdle models and their 95% credible intervals (CI) for activity, exploration and predation behaviours. The term VI refer to among individual variance in mean behaviour.

### 4. Effect of treatment on correlations between assays at the among individual level.

Table S3. Posterior correlation parameter estimates for among individual correlations in mean behaviour and associated 95% credible intervals (CI). Names of parameters starting with “Act”, “Exp” and “Pred” refer to activity, exploration and predation assays respectively. The term VI refers to among individual variance in mean behaviour.

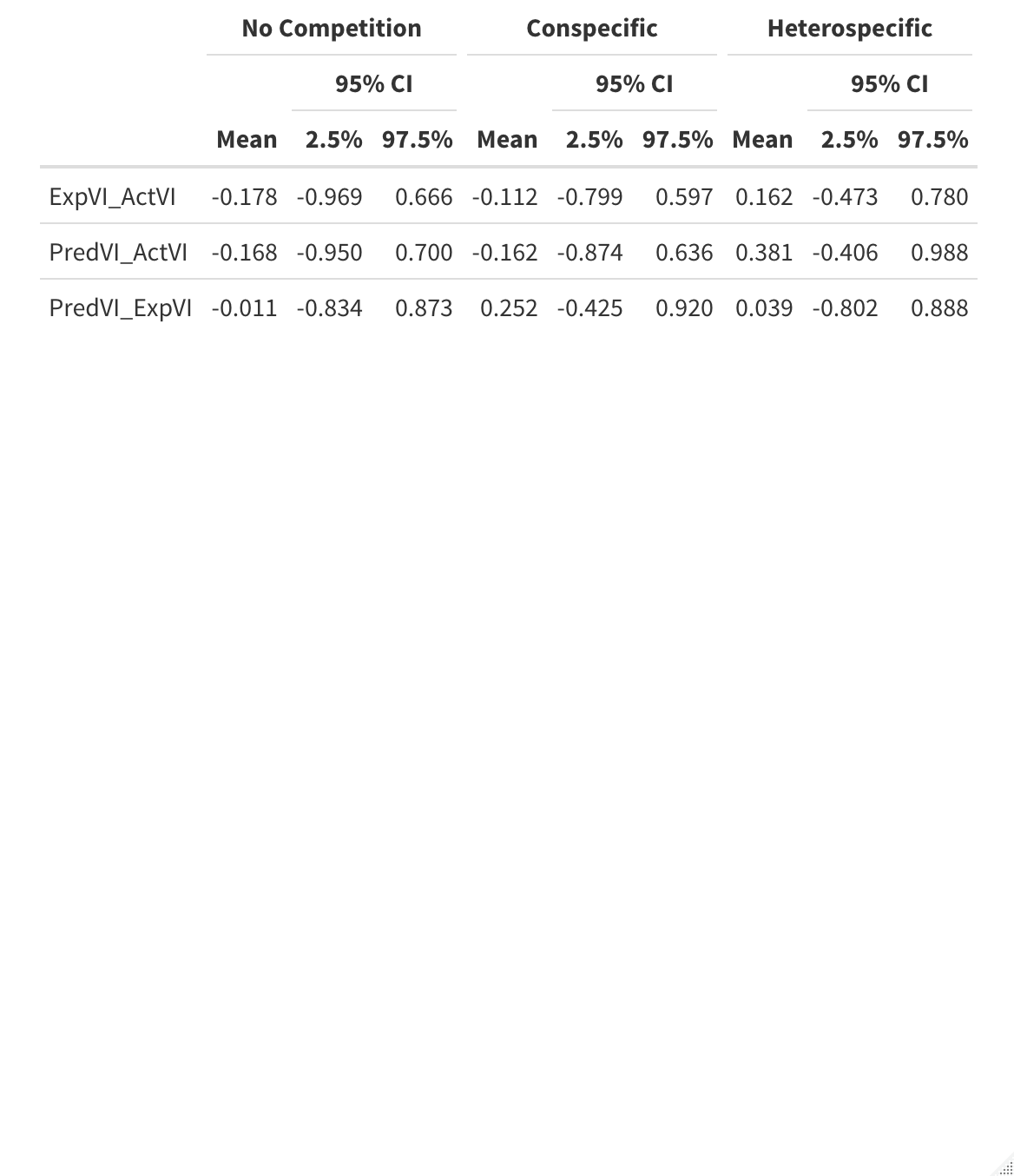

### 5. Treatment differences between assay correlations at the among individual level.

Table S4. Posterior correlation parameter estimates for the difference in among individual correlations in mean behaviour and associated 95% credible intervals (CI) for each treatment comparison. Names of parameters starting with “Act”, “Exp” and “Pred” refer to activity, exploration and predation assays respectively. The term VI refers to among individual variance in mean behaviour.

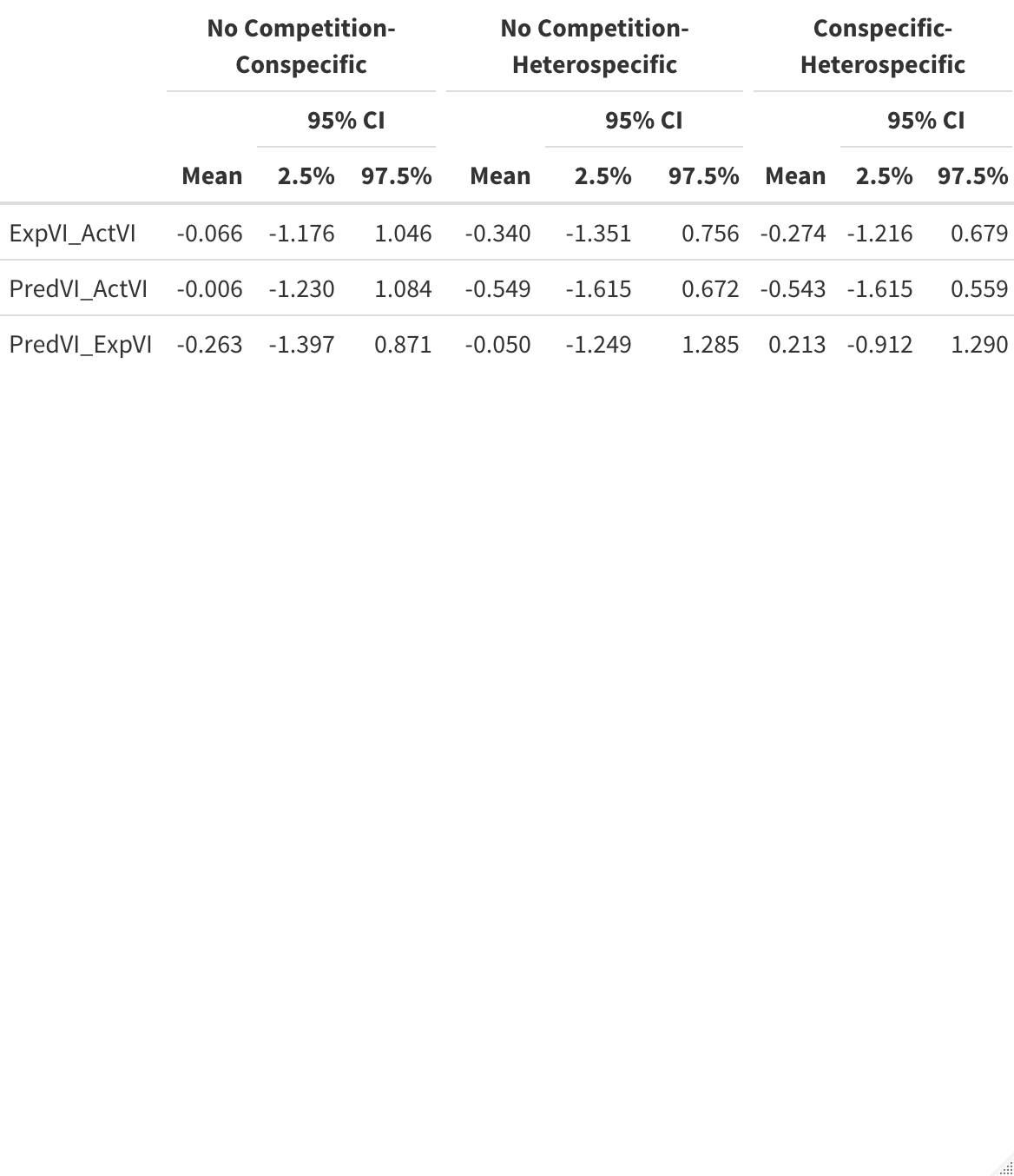
